# Key components of the delirium syndrome and mortality: greater impact of acute change and disorganised thinking in a prospective cohort study

**DOI:** 10.1101/251272

**Authors:** RA Diwell, D Davis, V Vickerstaff, EL Sampson

## Abstract

**Background:** Delirium increases the risk of mortality during an acute hospital admission. Full syndromal delirium (FSD) is associated with greatest risk and subsyndromal delirium (SSD) is associated with intermediate risk, compared to patients with no delirium - suggesting a dose-response relationship. It is not clear how individual diagnostic symptoms of delirium influence the association with mortality. Our objectives were to measure the prevalence of FSD and SSD, and assess the effect that FSD, SSD and individual symptoms of delirium (from the Confusion Assessment Method-short version (s-CAM)) have on mortality rates.

**Methods:** Exploratory analysis of a prospective cohort (aged ≥ 70 years) with acute (unplanned) medical admission (4/6/2007-4/11/2007). The outcome was mortality (data censored 6/10/2011). The principal exposures were FSD and SSD compared to no delirium (as measured by the CAM), along with individual delirium symptoms on the CAM. Cox regression was used to estimate the impact FSD and SSD and individual CAM items had on mortality.

**Results:** The cohort (n=610) mean age was 83 (SD 7); 59% were female. On admission, 11% had FSD and 33% had SSD. Of the key diagnostic symptoms for delirium, 17% acute onset, 19% inattention, 17% disorganised thinking and 17% altered level of consciousness. Unadjusted analysis found FSD had an increased hazard ratio (HR) of 2.31 (95%CI 1.71, 3.12), for SSD the HR was 1.26 (1.00, 1.59). Adjusted analysis remained significant for FSD (1.55 95%CI 1.10, 2.18) but nonsignificant for SSD (HR=0.92 95% CI 0.70, 1.19). Two CAM items were significantly associated with mortality following adjustment: acute onset and disorganised thinking.

**Conclusion:** We observed a dose-response relationship between mortality and delirium, FSD had the greatest risk and SSD having intermediate risk. The CAM items “acute onset” and “disorganised thinking” drove the associations observed. Clinically, this highlights the necessity of identifying individual symptoms of delirium.

## BACKGROUND

Delirium is an acute neuropsychiatric syndrome affecting around 25% of general hospital patients aged over 65 years [1-4]. It is characterised by acute onset and fluctuating course of disturbed attention, consciousness, orientation, memory, arousal and, behaviour, and alterations in perception and sleep cycle [5].

The aetiology of delirium is complex and multifactorial, including causes such as infection, sleep deprivation, pain, specific organ failures and metabolic disturbances [1, 6-8]. Each individual’s threshold for delirium differs depending on predisposing risk factors such as age and frailty [9].

Many operational definitions exist for delirium, including formal classifications in the Diagnostic and Statistical Manual of Mental Disorders (DSM) and algorithms such as the Confusion Assessment Method (CAM) [10]. Intermediate states, subsyndromal delirium (SSD), can be defined where individuals have symptoms of delirium but insufficient to meet the criteria for full syndromal delirium (FSD) [11].

FSD is associated with a number of poor outcomes, such as longer hospital stays, increased risk of post-hospital institutionalisation post-discharge, and accelerated cognitive decline [3, 8, 12-16]. FSD carries its own risk of death, independent of an individual’s exposure to established risk factors [3, 17-20]. The literature on SSD and adverse outcomes is less conclusive, partly because of variable definitions of SSD in relation to symptom clusters and/or severity [11, 21, 22].

It is possible that a dose-response relationship between FSD and mortality operates, such that SSD carries intermediate risk [23]. However, this has often not been systematically evaluated in the same cohort, using standardised definitions and maximally adjusting for a wide range of acute and chronic relationship observed. In particular, no studies have estimated mortality rates associated with individual diagnostic items from rating scales such as the CAM.

Our objectives were to: (1) examine the prevalence of FSD and SSD in a representative cohort of older acute hospital in participants over the age of 70 years; (2) estimate the impact of FSD and SSD (as measured by the short CAM (s-CAM) on admission) on mortality rates and (3) assess the impact individual key diagnostic items on the s-CAM have on this relationship.

## METHODS

### Design

We undertook an exploratory retrospective analysis of data collected on a cohort of older people with acute medical illness admitted into hospital between 4/6/2007 to 4/11/2007. Characteristics of the cohort have been described previously [24]. In brief, participants were eligible for inclusion if they were: ≥70 years old with an unplanned medical admission who were admitted >48 hours. All clinical assessments were conducted by psychiatrists within 72 hours of admission. Participants who lacked English language skills necessary to complete basic cognitive assessments were excluded. We sought verbal consent from participants or, if they lacked capacity to consent, verbal assent from their carers. The study involved the collection of routine clinical data that has subsequently been fully anonymised. The findings of these assessments were documented on the medical notes so that clinical teams could act on them if they wished. The exclusion of patients unable to give written informed consent or those without a relative to give assent for their participation may have caused selection bias, excluding the patient population we wished to study. The study and its verbal consent procedure was approved by the Royal Free Hospital NHS Trust Ethics Committee (06/Q0501/31).

### Outcome

Mortality was flagged by the UK Office for National Statistics (ONS) (mortality data censored 77 6/10/2011).

### Main exposures

#### Delirium

Participants were assessed using the Confusion Assessment Method, short version (s-CAM), which details the following delirium features: (1) acute onset, (2) inattention, (3) disorganised thinking, (4) altered level of consciousness [25]. The s-CAM has high sensitivity of >94% and specificity >90% for the detection of delirium and accurately distinguishes between delirium and dementia [26]. FSD was defined as persons demonstrating abnormalities in features 1 + 2 + (3 or 4). SSD was defined as having one or more s-CAM symptoms, but not fulfilling criteria for FSD. All participants without symptoms of FSD or SSD were defined as ‘no delirium’.

#### Covariates

Demographic data (age, sex, place of residence, ethnic origin and marital status) was collected from hospital records. Other assessments included the Charlson Co-morbidity Index [27, 28], Waterlow Scale [29] and a modified version of the Acute Physiology and Chronic Health Evaluation (APACHE II) [30-32](omitting the arterial blood gas). Severity of functional impairment prior to hospital admission was gathered from next of kin or other carers using the Functional Assessment Staging Scale (FAST) [33].

#### Data analysis

Differences in categorical and continuous variables according to delirium status were assessed using chi-square, ANOVA and Kruskal Wallis tests as appropriate. Continuous variables with skewed data (CCI and APACHE II scores) were categorised into standard quartiles for the final analysis.

Survival estimates for FSD, SSD and no delirium were compared using Kaplan-Meier curves and log-rank tests. Cox regression was used to examine the relationship between FSD, SSD and no delirium with mortality risk, sequentially adjusting for relevant confounders in a multivariable model. Finally, the relationship between each CAM criterion and mortality was estimated in the whole cohort, irrespective of syndromal status. Proportional hazard assumptions were met for all Cox regression analyses, confirmed by Schoenfeld Residuals ≥0.05. Candidate prediction models were compared using Harrell’s c statistics. Data were analysed using STATA version 12.

## RESULTS

### Study population

A total of 785 participants were recruited, of these, 75 participants had missing data and were excluded, leaving 710 participants assessed using the s-CAM at the time of admission. Exclusions occurred due to: incomplete/missing data (n=32, (5%), being too ill (n=18, (2%), untraceable (n=2, (1%), unable to speak English sufficiently (n=25, (3%), refusal to participate (n=23, (3%). Therefore, 610 (86%) participants from the original sample were included (Figure 1).

**Figure 1:**
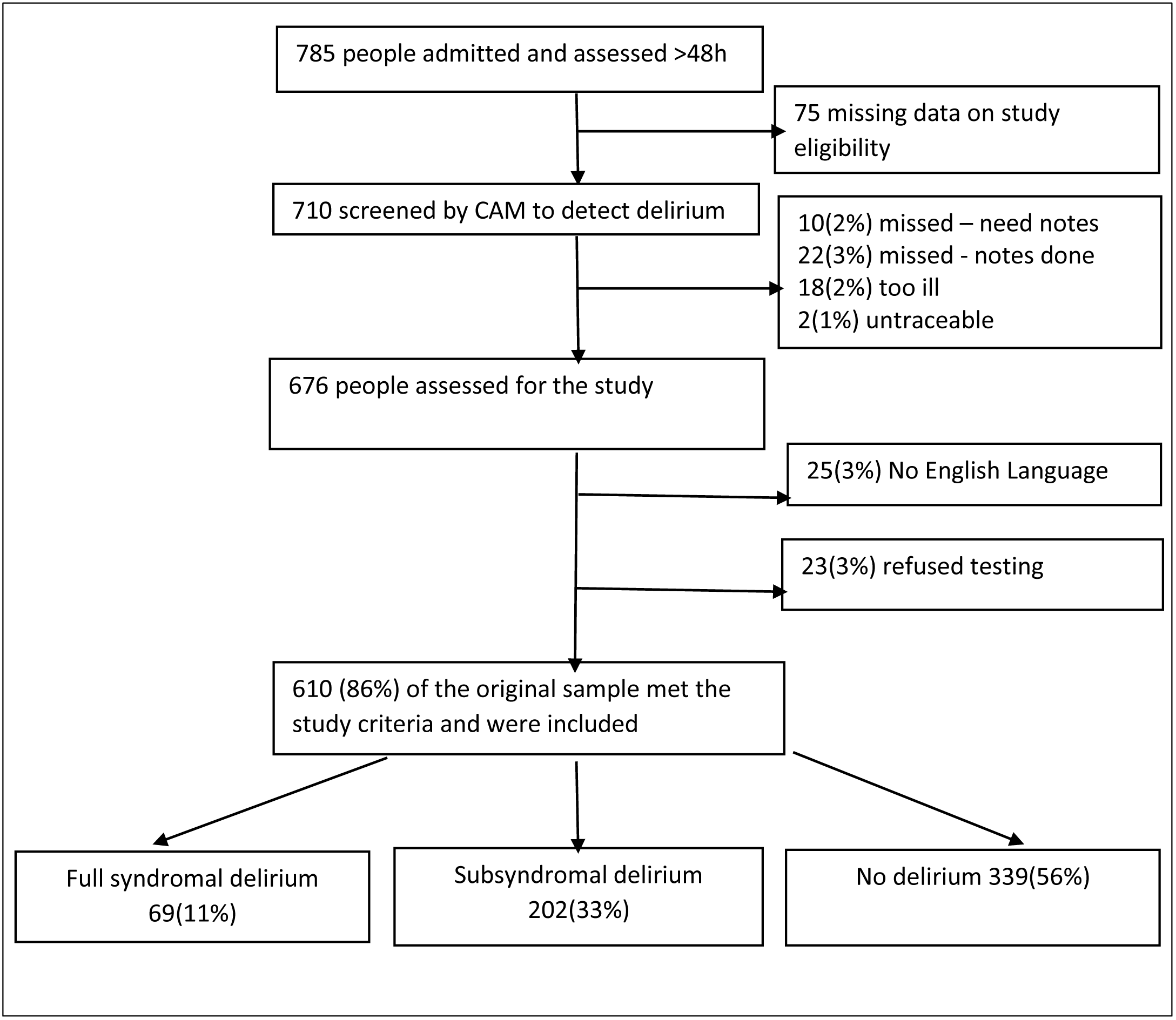
Study flowchart. Study flowchart showing the exclusion process and exclusion criteria for the study sample. 86% of the original sample were considered eligible for the study.

Mean age was 83 (sd 7) and over half were female (59%). A majority of the participants lived in their home (71%) and were of White British origin (70%) (Table 1).

**Table 1:**
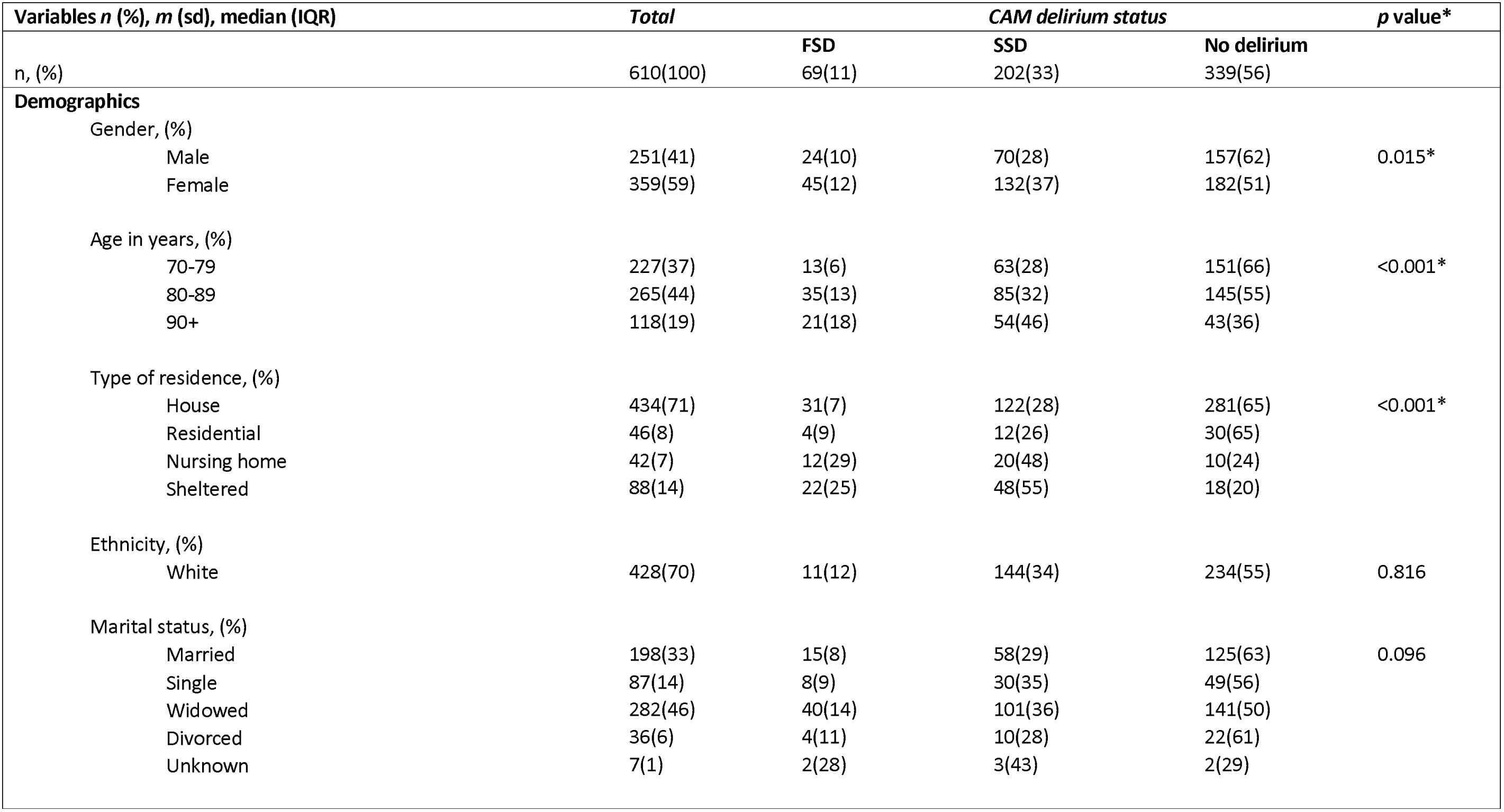
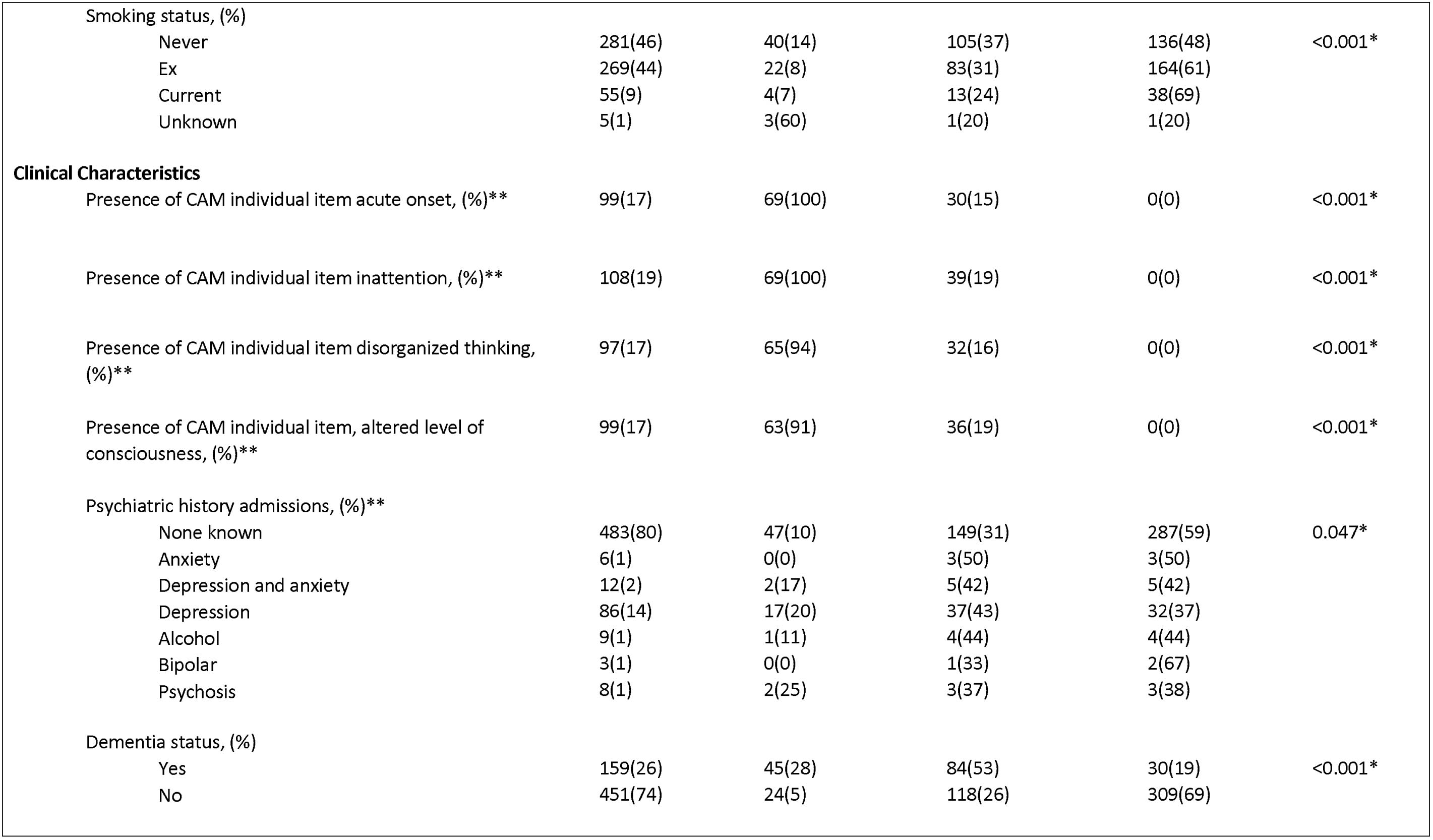
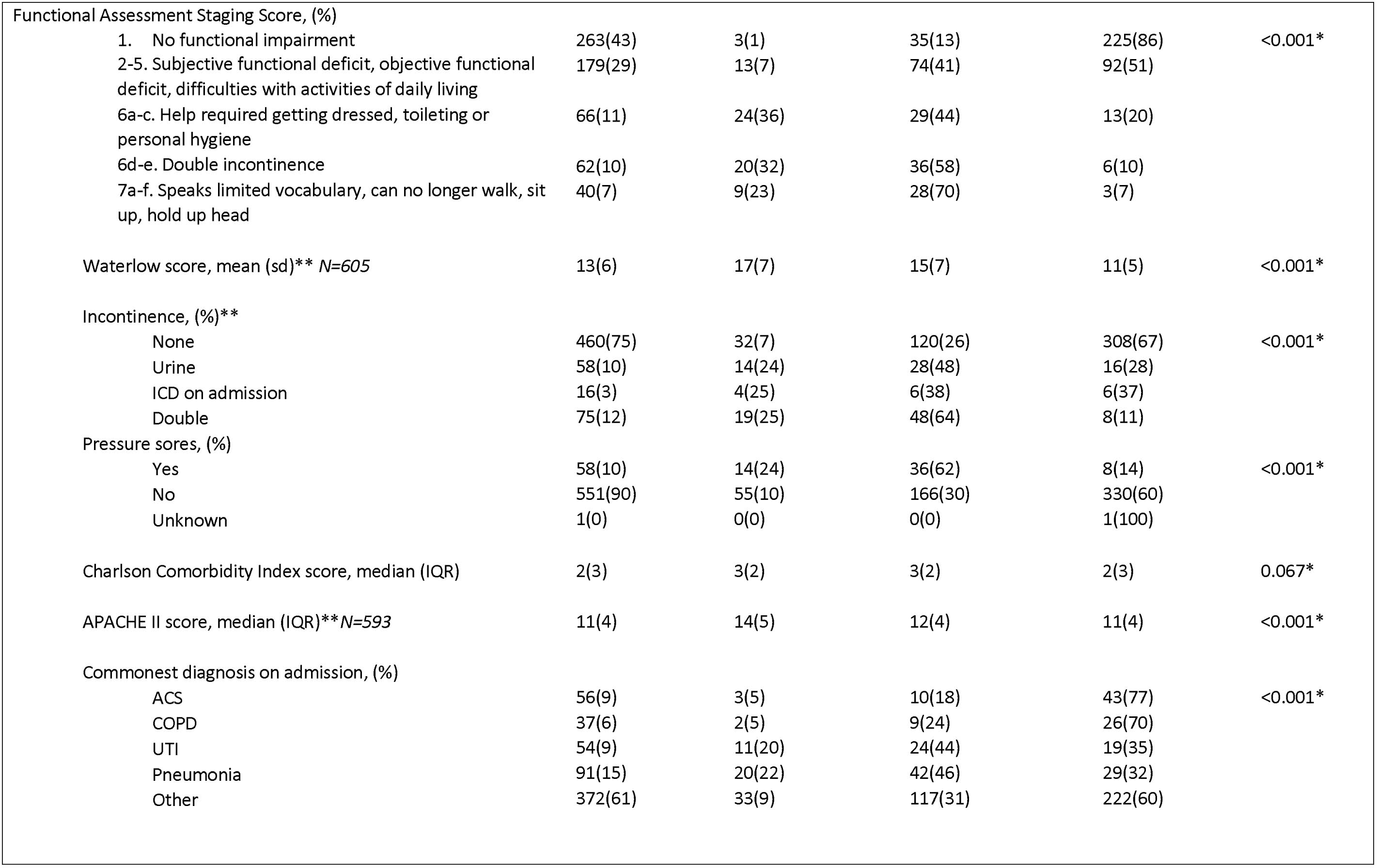
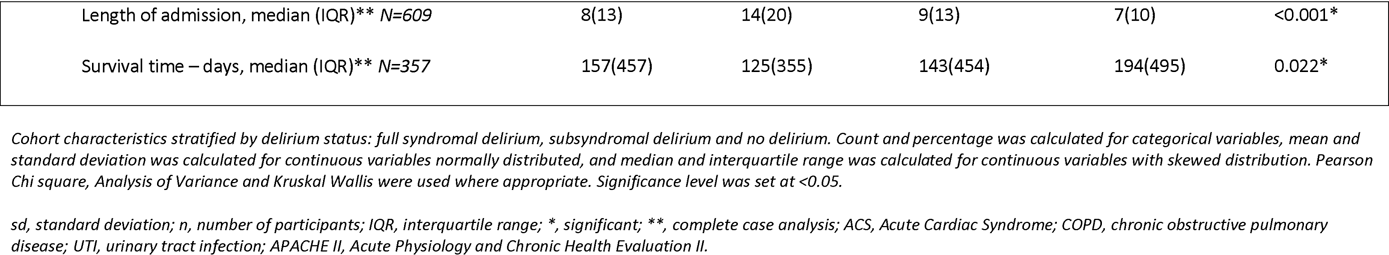
Cohort characteristics by CAM delirium diagnosis

| Variables *n* (%), *m* (sd), median (IQR) | Total | CAM delirium status |  |  | *p* value* |
| --- | --- | --- | --- | --- | --- |
| *n*, (%) | 610(100) | FSD<br>69(11) | SSD<br>202(33) | No delirium<br>339(56) |  |
| **Demographics** |  |  |  |  |  |
| Gender, (%) |  |  |  |  |  |
| Male | 251(41) | 24(10) | 70(28) | 157(62) | 0.015* |
| Female | 359(59) | 45(12) | 132(37) | 182(51) |  |
| Age in years, (%) |  |  |  |  |  |
| 70-79 | 227(37) | 13(6) | 63(28) | 151(66) | <0.001* |
| 80-89 | 265(44) | 35(13) | 85(32) | 145(55) |  |
| 90+ | 118(19) | 21(18) | 54(46) | 43(36) |  |
| Type of residence, (%) |  |  |  |  |  |
| House | 434(71) | 31(7) | 122(28) | 281(65) | <0.001* |
| Residential | 46(8) | 4(9) | 12(26) | 30(65) |  |
| Nursing home | 42(7) | 12(29) | 20(48) | 10(24) |  |
| Sheltered | 88(14) | 22(25) | 48(55) | 18(20) |  |
| Ethnicity, (%) |  |  |  |  |  |
| White | 428(70) | 11(12) | 144(34) | 234(55) | 0.816 |
| Marital status, (%) |  |  |  |  |  |
| Married | 198(33) | 15(8) | 58(29) | 125(63) | 0.096 |
| Single | 87(14) | 8(9) | 30(35) | 49(56) |  |
| Widowed | 282(46) | 40(14) | 101(36) | 141(50) |  |
| Divorced | 36(6) | 4(11) | 10(28) | 22(61) |  |
| Unknown | 7(1) | 2(28) | 3(43) | 2(29) |  |
| Smoking status, (%) |  |  |  |  |  |
| Never | 281(46) | 40(14) | 105(37) | 136(48) | <0.001* |
| Ex | 269(44) | 22(8) | 83(31) | 164(61) |  |
| Current | 55(9) | 4(7) | 13(24) | 38(69) |  |
| Unknown | 5(1) | 3(60) | 1(20) | 1(20) |  |
| Clinical Characteristics |  |  |  |  |  |
| Presence of CAM individual item acute onset, (%)** | 99(17) | 69(100) | 30(15) | 0(0) | <0.001* |
| Presence of CAM individual item inattention, (%)** | 108(19) | 69(100) | 39(19) | 0(0) | <0.001* |
| Presence of CAM individual item disorganized thinking, (%)** | 97(17) | 65(94) | 32(16) | 0(0) | <0.001* |
| Presence of CAM individual item, altered level of consciousness, (%)** | 99(17) | 63(91) | 36(19) | 0(0) | <0.001* |
| Psychiatric history admissions, (%)** |  |  |  |  |  |
| None known | 483(80) | 47(10) | 149(31) | 287(59) | 0.047* |
| Anxiety | 6(1) | 0(0) | 3(50) | 3(50) |  |
| Depression and anxiety | 12(2) | 2(17) | 5(42) | 5(42) |  |
| Depression | 86(14) | 17(20) | 37(43) | 32(37) |  |
| Alcohol | 9(1) | 1(11) | 4(44) | 4(44) |  |
| Bipolar | 3(1) | 0(0) | 1(33) | 2(67) |  |
| Psychosis | 8(1) | 2(25) | 3(37) | 3(38) |  |
| Dementia status, (%) |  |  |  |  |  |
| Yes | 159(26) | 45(28) | 84(53) | 30(19) | <0.001* |
| No | 451(74) | 24(5) | 118(26) | 309(69) |  |
| Functional Assessment Staging Score, (%) |  |  |  |  |  |
| 1. No functional impairment | 263(43) | 3(1) | 35(13) | 225(86) | <0.001* |
| 2-5. Subjective functional deficit, objective functional deficit, difficulties with activities of daily living | 179(29) | 13(7) | 74(41) | 92(51) |  |
| 6a-c. Help required getting dressed, toileting or personal hygiene | 66(11) | 24(36) | 29(44) | 13(20) |  |
| 6d-e. Double incontinence | 62(10) | 20(32) | 36(58) | 6(10) |  |
| 7a-f. Speaks limited vocabulary, can no longer walk, sit up, hold up head | 40(7) | 9(23) | 28(70) | 3(7) |  |
| Waterlow score, mean (sd)** N=605 | 13(6) | 17(7) | 15(7) | 11(5) | <0.001* |
| Incontinence, (%)** |  |  |  |  |  |
| None | 460(75) | 32(7) | 120(26) | 308(67) | <0.001* |
| Urine | 58(10) | 14(24) | 28(48) | 16(28) |  |
| ICD on admission | 16(3) | 4(25) | 6(38) | 6(37) |  |
| Double | 75(12) | 19(25) | 48(64) | 8(11) |  |
| Pressure sores, (%) |  |  |  |  |  |
| Yes | 58(10) | 14(24) | 36(62) | 8(14) | <0.001* |
| No | 551(90) | 55(10) | 166(30) | 330(60) |  |
| Unknown | 1(0) | 0(0) | 0(0) | 1(100) |  |
| Charlson Comorbidity Index score, median (IQR) | 2(3) | 3(2) | 3(2) | 2(3) | 0.067* |
| APACHE II score, median (IQR)**N=593 | 11(4) | 14(5) | 12(4) | 11(4) | <0.001* |
| Commonest diagnosis on admission, (%) |  |  |  |  |  |
| ACS | 56(9) | 3(5) | 10(18) | 43(77) | <0.001* |
| COPD | 37(6) | 2(5) | 9(24) | 26(70) |  |
| UTI | 54(9) | 11(20) | 24(44) | 19(35) |  |
| Pneumonia | 91(15) | 20(22) | 42(46) | 29(32) |  |
| Other | 372(61) | 33(9) | 117(31) | 222(60) |  |
| Length of admission, median (IQR)** N=609 | 8(13) | 14(20) | 9(13) | 7(10) | <0.001* |
| Survival time – days, median (IQR)** N=357 | 157(457) | 125(355) | 143(454) | 194(495) | 0.022* |

Cohort characteristics stratified by delirium status: full syndromal delirium, subsyndromal delirium and no delirium. Count and percentage was calculated for categorical variables, mean and standard deviation was calculated for continuous variables normally distributed, and median and interquartile range was calculated for continuous variables with skewed distribution. Pearson Chi square, Analysis of Variance and Kruskal Wallis were used where appropriate. Significance level was set at <0.05.
sd, standard deviation; n, number of participants; IQR, interquartile range; \*, significant; \*\*, complete case analysis; ACS, Acute Cardiac Syndrome; COPD, chronic obstructive pulmonary disease; UTI, urinary tract infection; APACHE II, Acute Physiology and Chronic Health Evaluation II.

A total of 69 (11%) participants had FSD, 202 (33%) had SSD and 339 (56%) had no delirium. The diagnostic symptom *inattention* had slightly higher prevalence (19%) compared to *acute onset, disorganised thinking and altered level of consciousness* (17%). Median CCI score was 2 (IQR 3) and APACHE II score was 11 (IQR 4), and the mean Waterlow score was 13 (6) (Table 1).

Prevalence of FSD and SSD increased with age, though there was no association with gender. FSD and SSD became more prevalent as age increased. Participants with FSD and SSD were more likely to live in nursing or sheltered accommodation. There was an overall higher prevalence of having a pre-existing dementia diagnosis, higher Waterlow scores, higher APACHE II scores and greater length of hospital stay.

Kaplan-Meier curves showed delirium was associated with reduced survival and that participants with FSD had greatest reduction in survival estimates compared to participants with no symptoms, and SSD had intermediate reduction (<0.001) (Figure 2). FSD had a median survival time of 5 months, compared to 21 months for SSD and 31 months for participants with no symptoms (Table 2).

**Figure 2:**
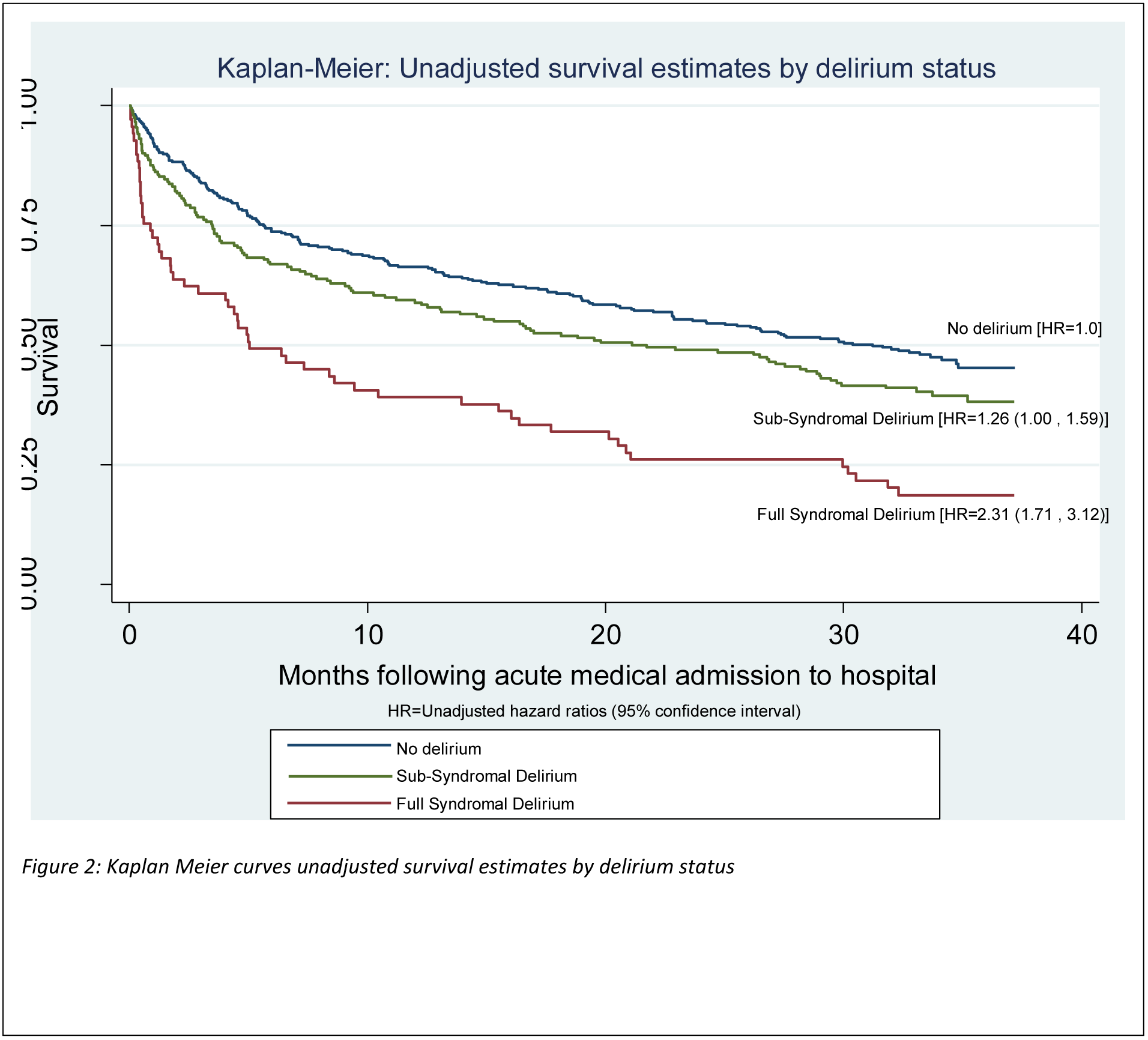
Kaplan-Meier: Unadjusted survival estimates by delirium status. Kaplan Meier curves illustrate unadjusted survival estimates by delirium status. Full syndromal delirium is shown to have significant reduction in survival estimates, compared to patients no symptoms. It also shows that subsyndromal delirium has intermediate reduction in survival estimates compared against full syndromal delirium and no symptoms.

**Table 2:** Mortality by delirium status (95%CI)

Delirium subtypes and mortality
| **Table 2: Mortality by delirium status (95%CI)** |  |  |  |
| --- | --- | --- | --- |
|  | **Delirium status** |  |  |
|  | *Full delirium n= 56* | *Subsyndromal delirium n=122* | *No delirium n=179* |
| *Survival time%* |  |  |  |
| <6 months | 62.50 (0.49 , 0.76) | 54.92 (0.46 , 0.64) | 49.72 (0.42 , 0.57) |
| >6 months | 37.50 (0.24 , 0.51) | 45.08 (0.36 , 0.54) | 50.28 (0.43 , 0.58) |
| *Median survival time (months)* | 5.03 (2.30 , 13.93) | 21.16 (13.11 , 29.04) | 31.21 (23.66 , NA) |

*Percentage of eligible patients and 95% confidence intervals stratified into survival time less than or more than 6 months following hospital admission. Death was flagged by the UK Office of National Statistics and certified by a death certificate. Median length and 95% confidence intervals for survival time was calculated following hospital admission. Complete case = 357.*
*<, less than; >, more than; NA, not available.*

In unadjusted Cox models, participants with FSD had a higher mortality risk (HR 2.31 95%CI 1.71, 3.12) compared with participants with no delirium. Participants with SSD had 1.26 (95% CI 1.00, 1.59) greater risk of mortality compared to participants with no symptoms. Each adjustment variable (age, gender, CCI, Waterlow and APACHE II) was independently related to death (p<0.001), except gender (p=0.684) (Table 4). Sequential adjustment showed that the associations between FSD and mortality remained after adjusting for age, sex, CCI, Waterlow and APACHE II (HR 1.55 95%CI 1.10, 2.18). The same sequence of adjustments for SSD and mortality showed greater attenuation (HR = 0.92 95% CI 0.70, 1.19). Unadjusted Cox models showed each s-CAM item was associated with higher mortality (p<0.001).

After sequential adjustment for age, sex, CCI, Waterlow and APACHE II, *acute onset* (HR 1.41 95% CI 1.07, 1.86) and *disorganised thinking* (HR 1.42 95% CI 1.05, 1.92) were associated with mortality, whereas this was no longer the case for estimates for *inattention* (HR 1.24 95% CI 0.92, 1.67) and *altered level of consciousness* (HR 1.33 95% CI 0.98, 1.79). C-statistics for all models were very close (0.66 to 0.67), suggesting comparable predictive ability for this set of variables.

**Table 3:** Adjusted cox regression model for the effect of the 4 core symptoms of delirium status on mortality, sequentially adjusted for clinically relevant covariates

| HR (95%CI) *p*-value. |  |  |  |  |  |  |
| --- | --- | --- | --- | --- | --- | --- |
|  | *None*<br>*(unadjusted)* | **+Age** | **+ Gender** | **+ CCI** | **+ Waterlow** | **+ APACHE II** |
| **Delirium key core symptoms** |  |  |  |  |  |  |
| Acute onset (*n*=583) | 1.88<br>(1.45 , 2.42)<br>*p*<0.001* | 1.80<br>(1.39 , 2.33)<br>*p*<0.001* | 1.80<br>(1.39 , 2.33)<br>*p*<0.001* | 1.76<br>(1.35 , 2.29)<br>*p*<0.001* | 1.46<br>(1.11 , 1.91)<br>*p*=0.007* | 1.41<br>(1.07 , 1.86)<br>*p*=0.016* |
| Inattention (*n*=576) | 1.80<br>(1.40 , 2.32)<br>*p*<0.001* | 1.74<br>(1.35 , 2.25)<br>*p*<0.001* | 1.75<br>(1.36 , 2.26)<br>*p*<0.001* | 1.73<br>(1.34 , 2.24)<br>*p*<0.001* | 1.33<br>(1.01 , 1.77)<br>*p*=0.044* | 1.24<br>(0.92 , 1.67)<br>*p*=0.152 |
| Disorganised thinking (*n*=563) | 2.06<br>(1.59 , 2.67)<br>*p*<0.001* | 1.97<br>(1.51 , 2.55)<br>*p*<0.001* | 2.01<br>(1.54 , 2.54)<br>*p*<0.001* | 1.94<br>(1.48 , 2.54)<br>*p*<0.001* | 1.52<br>(1.14 , 2.04)<br>*p*=0.005* | 1.42<br>(1.05 , 1.92)<br>*p*=0.024* |
| Altered level of consciousness (*n*=588) | 2.04<br>(1.58 , 2.63)<br>*p*<0.001* | 1.95<br>(1.50 , 2.52)<br>*p*<0.001* | 1.96<br>(1.51 , 2.53)<br>*p*<0.001* | 1.82<br>(1.40 , 2.37)<br>*p*<0.001* | 1.41<br>(1.06 , 1.88)<br>*p*=0.018* | 1.33<br>(0.98 , 1.79)<br>*p*=0.063 |

*Cox proportional hazard regression analysis for survival estimates for the four key core symptoms of full syndromal delirium. Unadjusted model complete case = 610. The same sample was used for the sequentially adjusted Cox proportional hazards regression model (age, gender, CCI, Waterlow and APACHE II). APACHE II and CCI scores were split into quartiles for the purpose of the analysis. There was no evidence of interactions, these, therefore were no longer considered. Proportional hazard assumptions were met, confirmed by Schoenfeld residuals $\geq 0.05$ . Significance level set at <0.05.*
*CAM, Confusion Assessment Method; HR, hazard ratios; CI, confidence intervals; *p*, significance level; N, number of participants; \*, significant; CCI, Charlson Comorbidity Index; APACHE II, Acute Physiology and Chronic Health Evaluation II.*

## DISCUSSION

We demonstrated a dose-response relationship between SSD, FSD and mortality, even after adjustment for a wide range of acute and chronic health factors. Individual s-CAM items contribute differentially to this relationship; *acute onset* and *disorganised thinking* appear to drive the association. Taken together, these findings emphasise that neuropsychiatric symptoms that arise in the context of acute illness in older people identified individuals at higher risk for dying.

This study had several strengths. The large cohort size and prospective data in a diverse socio-economic and ethnic population benefited from standardised assessments by experts and automatic notification of deaths from the UK Office of National Statistics. Data was collected within a 72 hour time-period after admission so it is not possible to establish whether cases of delirium were prevalent or incident and although the s-CAM has been shown to have good interrater reliability of 0.81-1.00 [34] we do not have data on this for our study. In keeping with other studies, limitations include the possibility of residual confounding. We identified FSD and SSD at a prevalence and associated with adverse outcomes consistent with the range established from systematic reviews [1, 2].

Participants with SSD had outcomes intermediate to those with no delirium and FSD – particularly in relation to acute illness severity, poor prognosis and outcomes, suggesting a dose-response relationship between delirium severity and mortality risk, which is in keeping with previous work [21, 23]. However, few other studies have been able to establish these associations while also accounting for a wide range of acute and chronic health factors.[35]

There is little literature exploring the individual mortality risk associated with each key symptom of delirium. We found each individual item on the short s-CAM was significantly associated with mortality, though *acute onset* and *disorganised thinking* had greater risk of mortality when all items were mutually adjusted.

A number of underlying mechanisms may explain the observed dose-response relationship between delirium and mortality. The causes of delirium can persist, which itself could lead to protracted delirium, prolonged hospital stays [17], and increased risk of death [36]. In turn, longer hospital stays could expose patients to a greater risk of iatrogenic harm [37, 38] for example: participants with hypoactive delirium have a greater risk of aspiration pneumonia, whereas participants with hyperactive delirium have greater risk of falls [39, 40] which in turn could cause longer hospital stays, further health deterioration and greater risk of death. Disorganised thinking could be a particularly adverse symptom because it may represent more profound neurocognitive disturbance particularly detrimental in frail, older participants predisposed to chronic and severe physical illness [3, 35, 41, 42].

### Conclusions

Emergency admission of an older patient presenting with FSD or SSD is a strong potential indicator of risk of death. Clinically it is important to be aware that each key symptom of FSD is strongly related to death, and participants presenting with just one symptom still carry an increased risk – highlighting the necessity of recognising each symptom separately. Better awareness of the mortality risk associated with delirium would strengthen arguments for early intervention, better treatment and quality of care, considering care plans and encouragement of discussion of prognosis with the patient and/or carer.

## LIST OF ABBREVIATIONS

ANOVA: analysis of variance
APACHE-11: Acute Physiology and Chronic Health Evaluation
CAM: Confusion Assessment Method
S-CAM: Short Confusion Assessment Method
CCI: Charlson Comorbidity Index
DSM: Diagnostic and Statistical Manual of Mental Disorders
FAST: Functional Assessment staging
FSD: full syndromal delirium
HR: hazard ratio
IQR: interquartile range
sd: standard deviation
ONS: Office for National Statistics
SSD: subsyndromal delirium

## DECLARATIONS

### Ethnical approval and consent to participation

The study was approved by the Royal Free Hospital NHS Ethics Committee (06/Q0501/31).

### Consent for publication

Not applicable.

### Availability of data and materials

The datasets used and/or analysed during the current study available from the corresponding author on reasonable request.

### Authors’ Contributions

ELS conducted the original study. RAD, VV and DD planned the data analysis. RAD analysed and interpreted data. ELS, DD and VV assisted with interpretation of the data outcomes. All authors contributed to the writing of the manuscript and approved the final manuscript.

## Acknowledgements

Not applicable

